# Predicting Treatment Outcomes from Adaptive Therapy — A New Mathematical Biomarker

**DOI:** 10.1101/2025.04.03.646615

**Authors:** Kit Gallagher, Maximilian A. R. Strobl, Philip K. Maini, Alexander R. A. Anderson

## Abstract

Standard-of-care cancer therapy regimens are characterized by continuous treatment at the maximum tolerated dose; however, this approach often fails on metastatic cancers due to the emergence of drug resistance. An evolution-based treatment paradigm known as ‘Adaptive Therapy’ has been proposed to counter this, dynamically adjusting treatment to control, rather than minimize, the tumor burden, thus suppressing the growth of treatment-resistant cell populations and hence delaying patient relapse. Promising clinical results in prostate cancer indicate the potential of adaptive treatment protocols, but demonstrate broad heterogeneity in patient response. This naturally leads to the question: why does this heterogeneity occur, and is a ‘one-size-fits-all’ protocol best for patients across this spectrum of responses?

Using a Lotka–Volterra representation of drug-sensitive and -resistant tumor populations’ dynamics, we obtain a predictive expression for the expected benefit from Adaptive Therapy and propose two new mathematical biomarkers (the Delta AT Score and the eTTP) that can identify the best responders in a clinical dataset after the first cycle of treatment. Based on prior theoretical analyses, we derive personalized and clinically-feasible optimal treatment strategies, based on individual patient’s tumor dynamics. These strategies vary significantly between patients, and so we present a framework to generate individual treatment schedules based on a patient’s response to the first treatment cycle. Finally, we develop metrics to identify which patients have the greatest sensitivity to unplanned schedule changes, such as delayed appointments, allowing clinicians to identify high-risk patients that need to be monitored more closely and potentially more frequently. Overall, the proposed strategies offer personalized treatment schedules that consistently outperform clinical standard-of-care protocols.

## 1 Introduction

Modern cancer treatment relies on a plethora of strategies, but these are traditionally focused on a singular, overarching goal: maximal cell killing [1, 2, 3]. While these approaches have led to remarkable increases in cancer survival times over recent decades [4], the emergence of treatment resistance often stands in the way of a complete cure [5]. In cases of cell resistance to treatment, these aggressive strategies subject the patient to unnecessary toxicity [6], and may even contribute to the emergence of resistance [7, 8]. Alternatively, strategies that modulate the drug dose [9], or prescribe intermittent drug holidays [10], have been proposed to control the tumor, instead of trying to eliminate it. These strategies aim to make better use of existing drugs, extending the period over which a tumor is sensitive (and responsive) to a therapeutic agent.

One of the most promising approaches uses treatment holidays based on the principle of ‘competitive suppression’ - an evolutionary principle that drug-sensitive and -resistant subclones will compete, reducing the proliferation rate of resistant subclones and enabling tumor control. This is often supported by a ‘cost of resistance’, a reduction in the base proliferation rate of resistant cells relative to sensitive cells in the absence of treatment, which has been observed experimentally *in vitro* [11] and *in vivo* [12, 13] for several tumor cell lines.

In contrast to intermittent treatment strategies, which apply drug holidays of a standardized duration [14, 15], adaptive therapy (AT) modifies either the drug dosage or the holiday timing according to the patient’s disease dynamics [16]. In a pilot trial by Zhang et al. [17], drug holidays were scheduled according to the current tumor size: treatment was applied until the tumor burden halved (as measured by prostate-specific antigen biomarker (PSA) - a proxy for tumor burden), and then reapplied once it regained the original size. This led to significant inter- and intra-patient variation in holiday duration; for example, the average holiday duration ranged from 53 to 347 days between patients. Overall, patients undergoing AT had significantly improved median time to progression (TTP; 33.5 months) compared to 14.3 months in the standard-of-care cohort [18].

While demonstrating the potential of AT, this seminal work by Zhang et al. has provoked further questions - firstly whether the 50% threshold size used was optimal, and indeed whether all patient schedules should be determined by the same threshold size? Furthermore, these authors [18] observed significant variation in AT response within their cohort, with four patients in the study cohort still undergoing stable treatment cycles 53–70 months after the start of treatment. Is it possible to predict the benefit a patient will experience from AT, and could we identify these ‘super-responders’ in advance?

### 1.1 Designing Adaptive Therapy Schedules

Previous work in adaptive treatment schedules has demonstrated that the TTP may be maximized by maintaining as large a sensitive sub-population as possible, to maximize competitive suppression of the resistant subpopulation [19, 20]. We demonstrate this in Figure 1a, evaluating different treatment schedules on a virtual patient model (defined in Section 4.1), with parameter values taken from Strobl et al. [21] and given in Table 1. Specifically, we evaluate these schedules on a tumor profile with zero cost of resistance and zero cell turnover, shown to progress fastest under continuous therapy (CT) and benefit least from AT [21]. This demonstrates the benefit of AT on tumors that fail fastest under CT, currently the standard of care for most therapeutic agents. In this context we define treatment failure, or tumor progression, as a 20 % growth from the initial size (i.e. 1.2*N*_0_), following prior studies in this area (e.g., [11, 21, 20]).

**Table 1:** Parameter values/ranges used for the virtual patient.

| Name | Description | Value/Range | Reference |
| --- | --- | --- | --- |
| $r_S$ | Sensitive cell proliferation rate | $0.027 \text{ day}^{-1}$ | Adopted from [17] |
| $r_R$ | Resistant cell proliferation rate | $0.5r_S - 1.0r_S$ | Lower limit [11], upper limit of no cost |
| $d_S, d_R$ | Natural cell death rate | $0.0r_S - 0.5r_S$ | Lower limit given by zero turnover<br>Upper limit adopted from [38] |
| $d_D$ | Drug-induced cell killing | 1.5 | Adopted from [39] |
| $K$ | Tumor carrying capacity | 1.5 – 5 | Values in this range reported by [40] |
| $N_0$ | Initial tumor cell density | 0.1 – 0.75 | Rescaled by initial PSA measurement |
| $R_0$ | Initial resistant cell fraction | $0.0001N_0 - 0.1N_0$ | Values in this range reported by [41] |

**Figure 1:**
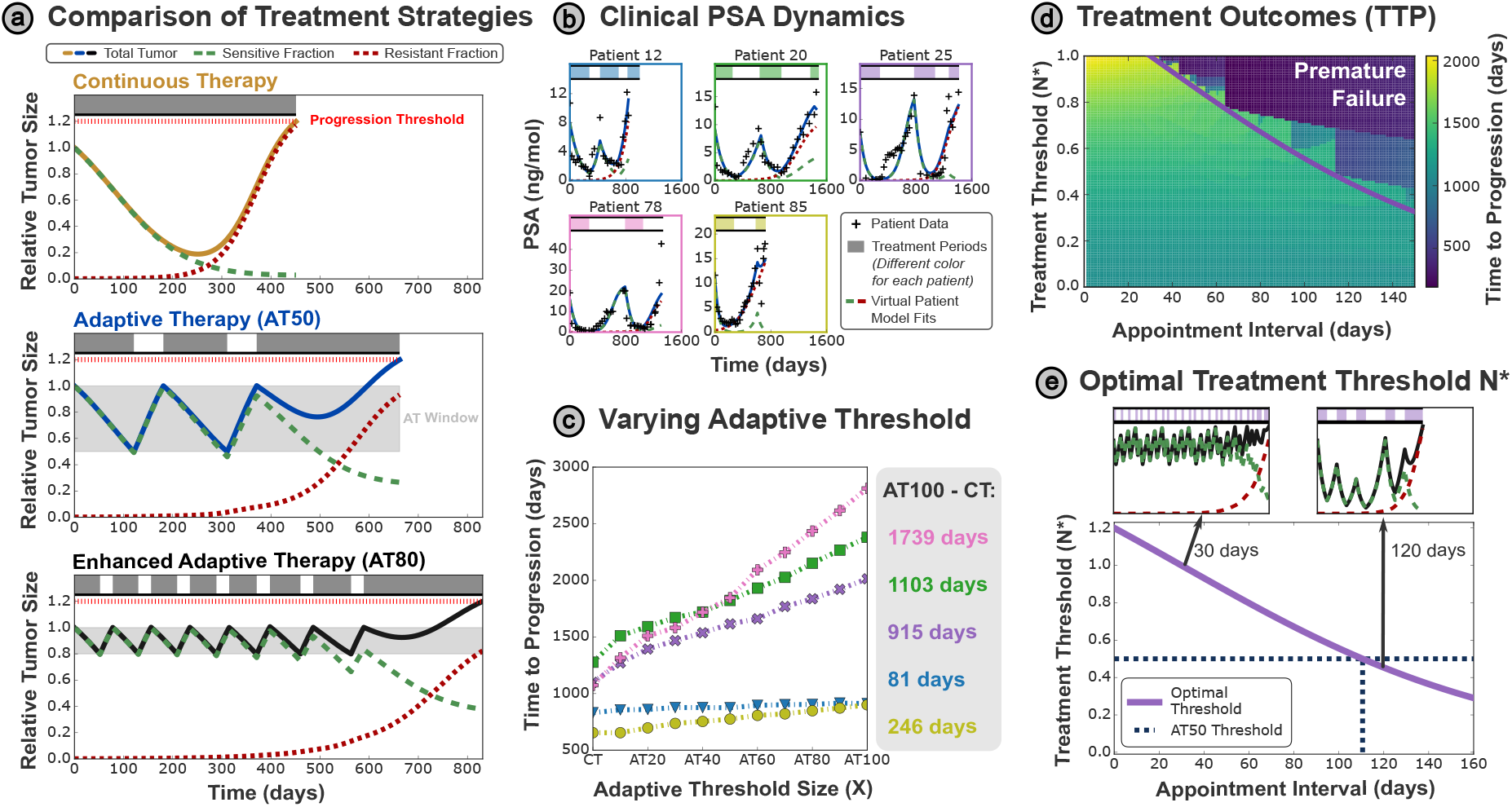
**(a)** Comparing treatment schedules, a conventional adaptive schedule (AT50) using a 50% size threshold (maintaining the total tumor size in a window between the initial tumor size *N*_0_ and 0.5*N*_0_, where treatment periods are denoted by shaded bars at the top of the figure) outperforms continuous therapy. Raising this threshold to 80% (i.e. AT80) provides further benefits, through increased suppression of the resistant population. **(b)** To explore personalization of the treatment threshold, we selected five patient profiles from Bruchovsky et al. [25], to which we fitted a Lotka-Volterra ordinary differential equation (ODE) model to form the virtual patient cohort used throughout this paper. **(c)** By raising the tumor size threshold required for a drug holiday to be initiated, we retain a larger sensitive population which generates greater competitive suppression of the resistant cells. This results in significant simulated increases of TTP for all of the patients in (b), though the maximal possible benefit of AT (given by the difference in TTP between AT100 and CT) varies significantly between patients (each given in a different color). **(d)** The analytically optimal threshold *N* ^∗^ can be plotted over the space of TTP outcomes - the TTP is maximized at higher treatment thresholds, however these are only possible for sufficiently frequent appointments. Treatment according to threshold values below *N* ^∗^ (for a given appointment spacing) will be sub-optimal, while thresholds above *N* ^∗^ may result in early tumor progression. **(e)** The curve *N* ^∗^(*τ*) varies strongly with the appointment interval *τ* (see equation (4), meaning that AT50 is only optimal for a specific appointment interval (~ 110 days). Intervals shorter than this benefit from a higher threshold tumor size, while longer intervals require a smaller threshold. Examples of this are shown in the insets for 30 and 120 days - note that treatment periods under the shorter interval are much more regular.

We find that the AT implemented by Zhang et al. [17], wherein treatment is applied until the tumor halves in size (AT50; defined in Equation 3), can gain an expected increase in TTP of more than 200 days over CT (shown in Figure 1a). However considering an alternate AT implementation (AT80 - wherein treatment is only applied until the tumor shrinks to 80% of its original size) shows further gains in TTP. The larger average tumor size in AT80 exerts a greater competitive suppression effect on the drug-resistant population, further delaying the tumor progression.

A natural extension of this trend has been suggested by Viossat and Noble [20], where retaining a large but tolerable tumor size is preferable for extending the TTP. Despite a range of theoretical work in this area [19, 22, 23], previous approaches rely on continuous monitoring of the disease and instantaneous treatment re-evaluation when the tumor size crosses a pre-defined threshold. However, clinical decisions are based on tumor information obtained at discrete time points, potentially adding a significant delay before changes in tumor dynamics are detected.

This practical reality limits our ability to maximize the benefit of AT, which necessitates frequent interventions to retain a large sensitive subpopulation (note the increased frequency of treatment holidays for AT80 in Figure 1a). We have previously shown [24] that there is a tradeoff between the interval between clinical appointments (where the tumor size is measured and treatment may be approximated) and the threshold tumor size that may be used for AT. More frequent appointments allow greater control over the tumor so that it may be maintained at a higher average size, and ultimately a longer TTP. However, this desire to treat frequently must be balanced with constraints on clinical resources and patient quality of life; it is simply infeasible to bring the patient into the clinic every day to reevaluate their treatment schedule.

In this paper, we aim to explore the practical implementation of optimal adaptive therapy in the clinic. We first demonstrate the heterogeneity in AT response between patients, also observed in previous clinical trials [18]. This results in subsequent heterogeneity in the optimal treatment schedule, with different patients benefiting most from different AT protocols based on their tumor dynamics.

In a clinical context, we would have no knowledge of these tumor dynamics for a new patient at the start of their treatment. We therefore propose a probing cycle - a single treatment cycle of AT50 - to characterize the drug response over single treatment cycle. We then use information from this to predict which patients will benefit the most from adaptive therapy, what optimal threshold a patient should have, and how often they should be seen (their appointment frequency). We propose two novel mathematical biomarkers - the Delta AT score which identifies patients that are expected to benefit most from AT, and the eTTP to quantify the absolute TTP extension that AT might confer for these patients. This approach provides a practical and accessible biomarkers to develop treatment strategies tailored to the individual patient

Finally, we consider possible changes to the clinical appointment frequency, exploring scenarios where clinicians must prioritize which patients to see more frequently. We present a prospective case study implementing these approaches on a cohort of exemplar virtual patients. Through this, we demonstrate how analytic tools can motivate changes to AT strategy through personalization of both the treatment threshold and appointment frequency, and identify those patients who will benefit the most from more frequent treatment interventions.

## 2 Results

### 2.1 Optimizing AT with Discrete Appointment Intervals

To explore the impact of varying the treatment threshold, we simulate virtual patient profiles taken from a Phase II intermittent therapy trial conducted by Bruchovsky et al. [25] (see Section 4.5 for more details). These patients were selected from the clinical trial to highlight the possible benefits enabled by personalized AT, since they all progressed and have distinct dynamics. Figure 1b shows the patient PSA records and demonstrates that the virtual patient model provides a good representation of the observed dynamics.

Simulating different AT protocols while varying the threshold *X* at which drug holidays are initiated (ATX protocol from Section 4.2, Figure 1c) shows that all patients have a higher TTP under AT than under CT. Furthermore, this benefit consistently increases as the threshold is raised, since larger sensitive populations are retained throughout treatment. However, the extent of this benefit varies significantly between patients, highlighted by the absolute difference in TTP between AT100 (the best performing AT schedule) and CT recorded for each patient in Figure 1c. AT100 delays progression for Patient 78 by almost five years, whereas the difference for Patient 12 is less than three months, highlighting the need for a predictive biomarker to identify which patients would benefit most from AT in advance of treatment.

#### 2.1.1 Computing a Personalized Treatment Threshold

When AT is implemented clinically, changes to the treatment regimen are based on PSA measurements from routine blood tests, and implemented through consultation with a clinician. For outpatients, their treatment plans may only be updated during regular clinical appointments, which may occur at most every two months. The discrete time interval between appointments poses an important practical challenge to AT: the frequency of clinical intervention required for the optimal AT100 strategy may not be practical to implement for many patients.

To address this issue, we propose to use mathematical modeling to personalize the AT threshold to balance the benefits from increased competitive suppression with the risks of premature progression given the patient’s appointment frequency and treatment dynamics. We will then recommend that patients receive treatment while the tumor is larger than this threshold size (which we will call *N* ^∗^), and only receive a treatment holiday when the tumor size is below *N* ^∗^.

We simulated the relationship between TTP, threshold size and appointment frequency for Patient 25 in Figure 1d, emphasizing the trend that a higher threshold tumor size results in a greater TTP. However, this only applies when the treatment plan is re-evaluated sufficiently frequently - treatment protocols that have insufficiently frequent clinical appointments for a given threshold tumor size are inherently risky, resulting in the variable (and often poorer) treatment outcomes observed to the right of Figure 1d.

An analytic expression for *N* ^∗^, defining the optimal threshold size for AT with discrete intervals of a given duration between appointments, is given in Section 4.3). Superimposing this relationship on the treatment outcomes for Patient 25 in Figure 1d, we can see that it effectively separates out the possible treatment protocols where the AT protocols that effectively extend the TTP (above the curve) from the combinations of *N* ^∗^ and *τ* that result in early tumor progression.

Beyond this, we see in Figure 1e that higher optimal thresholds necessitate increasingly frequent clinical appointments, up to the analytically optimal threshold of 1.2*N*_0_ which would require the treatment plan to be re-evaluated continuously. This trend is exemplified in the inset panels, where a shorter appointment interval (of 30 days) allows better control over the tumor size, keeping it at a more consistent, and higher average, size than a 120 day interval. The larger pool of sensitive cells results in more effective competitive suppression of the drug-resistant cells, and hence an extended TTP.

### 2.2 Application of a Probing Cycle

However, discussions of a personalized treatment threshold are futile without knowledge of patient-specific tumor dynamics and treatment response to base it on. This is much more limited in a clinical setting, when a new patient is presented for the first time and has had no previous exposure to this treatment. How can we efficiently collect sufficient information about the patient’s tumor dynamics to enable prediction of these personalized treatment metrics such as *N* ^∗^?

To address this issue, we have previously proposed delivery of a single AT50 cycle to ‘probe’ the patient’s treatment dynamics [26]. By collecting regular data on the tumor burden during this cycle, we can fit our mathematical model (1) to the probing cycle data (Figure 2a), identifying key model parameters primarily associated with the drug response and tumor regrowth rate under drug holidays. These can then be used to predict the benefit a patient may expect from AT, as well as determining the optimal adaptive strategy for them. Similar initial treatment cycles have previously been employed to compare patient response to different drugs, and identify potential collateral sensitivities [27]. This dynamic approach of re-evaluating treatment schedules based on observations of the patient response has been proposed previously to personalize treatment strategies to each patient’s tumor state [28].

**Figure 2:**
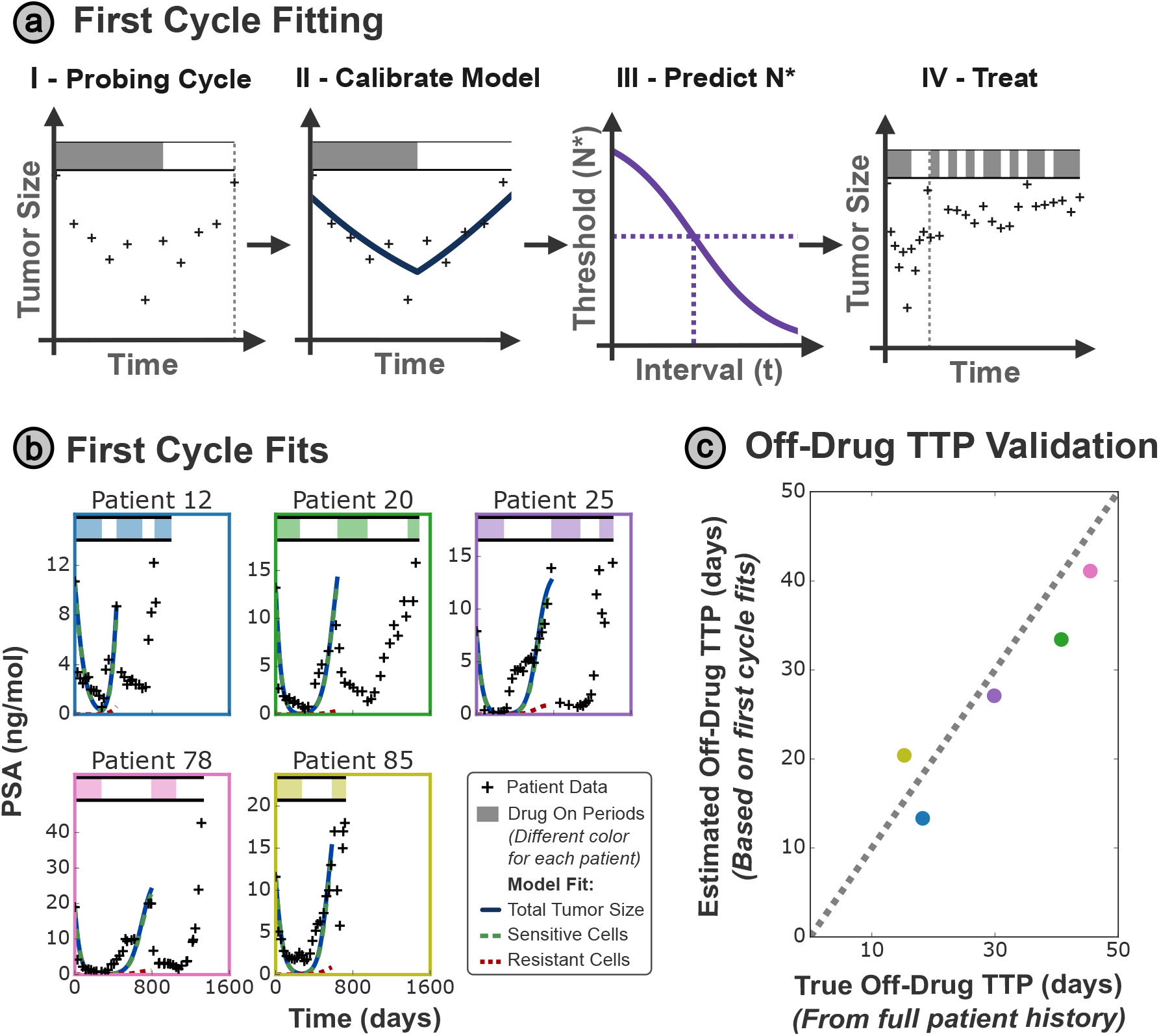
**(a)** We propose a probing cycle to predict the individual benefit from adaptive treatment schedules. Data from a standardized initial cycle of AT50 are fitted to the virtual tumor model, allowing prediction of the benefit of AT and personalization of the treatment strategy in future drug cycles. **(b)** Partial fits of the initial cycle of each patient’s clinical data accurately capture the heterogeneity in treatment response and tumor regrowth rates. **(c)** The parameter values derived from the probing cycle fits then accurately capture tumor dynamics, such as the off-drug TTP, demonstrating good agreement with the parameter values derived from the full patient history. The dashed line has a gradient of unity, and represents a perfect prediction.

To illustrate this, we generated model fits on clinical data from the initial treatment cycle for the 5 patients in our test cohort (Figure 2b). We separately fit each patient’s full treatment response to (1) to create a virtual patient profile, which can simulate the response to different treatment schedules for in silico validation of these predictions. This process is described in full in Appendix A.1.1. To test this fitting approach is able to capture the key dynamics of the tumor, we calculate the time taken for a tumor to grow 20% in the absence of drug (i.e. an off-drug equivalent of the TTP) for the estimated parameter set (from the probing cycle), and compare this to the simulated behavior of the virtual patient (determined by the full treatment history of the patient). In Figure 2c, we show that the probing cycle parameters are able to accurately replicate the relative ordering and exact values of the drug-free progression times within this test cohort, exemplifying the potential of this method to estimate key tumor dynamics.

#### 2.2.1 The Delta AT Score - Predicting the Relative Benefit of AT

Previous clinical trials have shown mixed responses to adaptive treatment schedules [18], and it has been verified computationally that some patients will benefit more than others even under treatment schedules close to optimal [26]. This begs the question: could we use these probing cycle fits to predict which patients would benefit most from AT?

AT is designed to utilize the competitive suppression of the resistant sub-population to delay progression. We therefore look to estimate the strength of competitive suppression from the probing cycle. Through this, we derive two metrics the Delta AT score and the eTTP, which predict the relative benefit from AT, and the expected TTP under any given adaptive strategy, respectively.

The ‘*Delta AT score*’ (Δ), which quantifies the relative benefit in TTP of AT (over CT) expected for a given patient, based on parameters measured in the probing cycle. The Delta AT score is based on the reduction in drug-resistant cell growth expected due to the presence of a sensitive population within the tumor. We expect that patients with a larger Delta AT score will experience a greater increase in their TTP on AT compared to their TTP for CT. Note that the Delta AT score is not designed to give an absolute prediction of extension in TTP that AT will provide (later we present the eTTP to provide this), but rather a relative score to indicate which patients will benefit most from AT.

To test the prognostic performance of the Delta AT Score we performed a simulation study. We calculated Δ using the parameter values estimated from the probing cycle (methods as in Appendix A.1.2) and benchmarked the score against the simulated benefit of AT over CT for the virtual patient counterparts. We find that the Delta AT score (based only on data from the probing cycle) is an excellent predictor of the observed benefit of AT, predicting the relative differences in simulated benefit between patients with an *R*^2^ = 0.92. Furthermore, the accuracy of the Delta AT score is replicated when predicting the benefit of simulated AT50 over the intermittent protocol received by the patients in the clinical trial, correctly ordering the patients by measured benefit with an *R*^2^ = 0.97. Most critically for clinical decision-making, this metric provides a relative ordering between the patients, representing who will benefit most from AT instead of quantifying the absolute extent of this benefit.

#### 2.2.2 The eTTP - Predicting the Absolute Benefit of AT

By combining the Delta score with further analytical results obtained from the probing cycle, we can also derive a second metric - the estimated TTP (eTTP), which provides an absolute estimate of the time-span by which AT may delay progression (detailed in Section 4.4.2).

Using the expected TTP for a fully-resistant tumor (in the absence of any sensitive cells) as a baseline, we scale by the Delta AT score to give an estimate for the TTP in the presence of a fixed sensitive population. This is visualized in Figure 3a, where panel I shows the TTP in absence of any sensitive cells, and panel II depicts how the TTP is extended by a factor of Δ due to the presence of a fixed population of sensitive cells, which slow the resistant growth dynamics. The size of this fixed population is determined by the adaptive strategy chosen; here we demonstrate an ATX strategy where the tumor is confined between two threshold sizes, and the average size is given by the arithmetic mean of the limits of this size window. In clinically realistic treatment strategies, the tumor size will overshoot this target window; accounting for this we obtain a patient-specific estimated TTP (eTTP) score.

**Figure 3:**
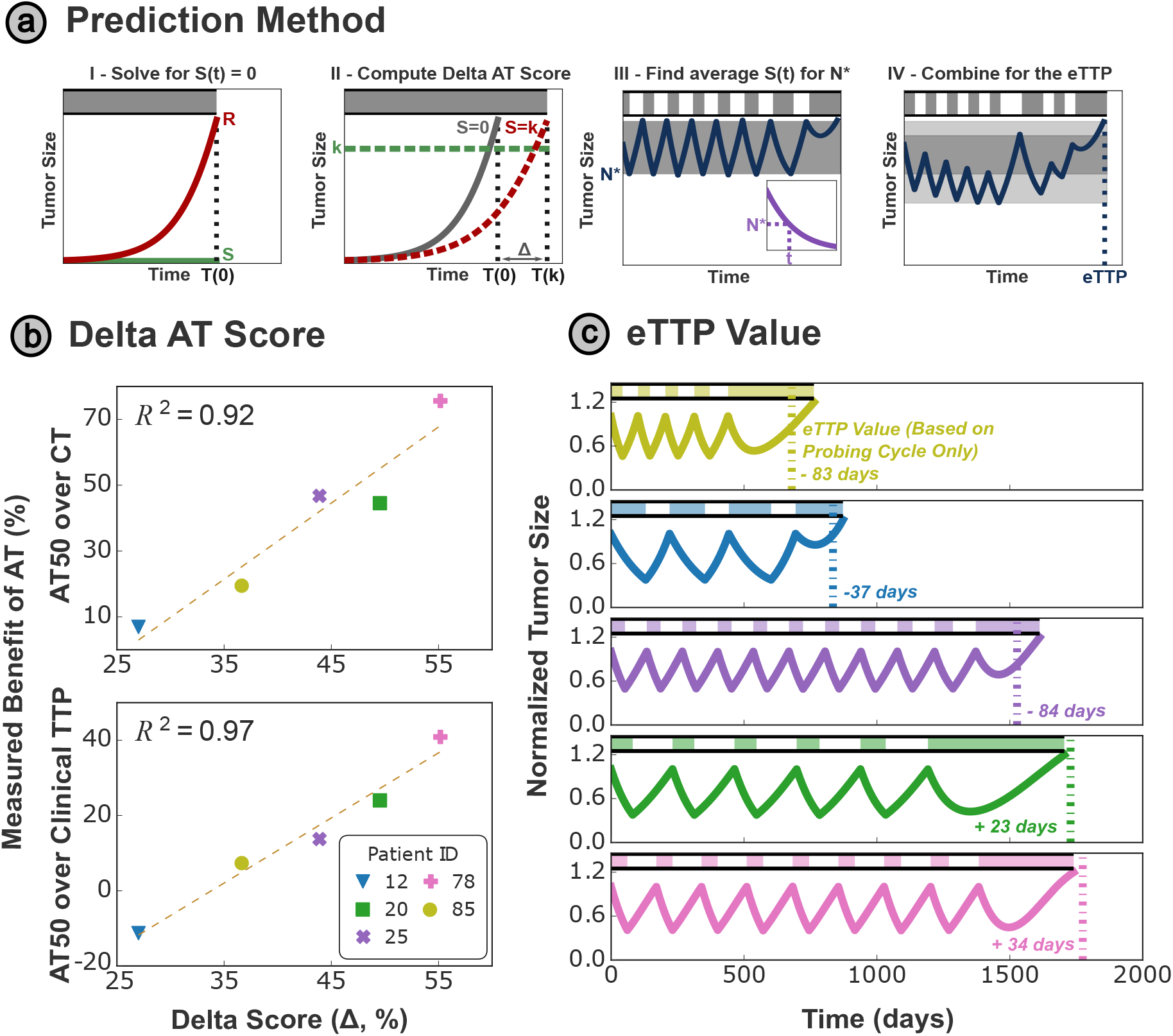
**(a)** We may estimate both the TTP, and the expected benefit from AT, by considering the simpler case of a fully resistant tumor, which we can solve analytically (I) to obtain a base-line (worst case) TTP: *T* (0). We then compute the benefit of AT conferred by a fixed sensitive population (II), which suppresses the growth of the resistant cells and proportionally increases the TTP to *T* (*κ*). Finally, we convert to the TTP benefit using the average tumor size (III - plotted in blue), assuming that the population will be constrained between the progression limit (20 % growth from the initial tumor size - 1.2*N*_0_) and the chosen treatment threshold *N* ^∗^ (based on the appointment interval *τ*). We finally correct this average size for overshooting (IV), where the tumor size does not reach the top gray shaded region, and overshoots past the lower threshold into the lower light gray region instead. **(b)** The predicted benefit metric (based on parameter values from the probing cycle alone) is strongly correlated with the observed benefit of AT50 - the increase in TTP observed for simulated AT50 (based on parameter values from the patients’ full history) compared to both simulated CT, and the clinical outcomes observed for this patient cohort. *R*^2^ values listed are from a linear regression, with the associated best fit line plotted in dashed gray. **(c)** Extending these predictions to the full TTP under AT50, we compare the eTTP (dashed lines) to simulated data based on the full-history parameters (solid lines), and replicate both the relative progression order and absolute TTP for each patient within the test cohort with a high degree of accuracy. (Errors are highlighted in days in the insets).

In Figure 3c we test the accuracy of the eTTP on the virtual patient cohort by comparing the prediction with the simulated TTP for AT50. The relative ranking of the TTP between patients is replicated exactly, with an average absolute error of 4.8% on the magnitude of the TTP, confirming the potential of this approach to predict treatment outcomes at the start of treatment. There is marginally more inaccuracy in these predictions than the Delta AT score depicted in Figure 3b - it is much more challenging to predict the absolute benefit of AT due to the reliance of this approach on parameters such as the initial number of resistant cells (*R*_0_). This parameter cannot be estimated from the probing cycle, as the dynamics of the small resistant fraction are masked by the much larger sensitive fraction, and so must be replaced by population-wide estimates instead (Appendix A.1.2). However, the eTTP offers a complementary perspective to the Delta AT score, quantifying the number of days by which AT may be expected to delay progression, helping inform both patients and clinicians of the benefit that AT might be expected to offer.

While it might seem intuitive to simply estimate the TTP via a computational simulation of AT50 once estimates of the tumor parameters have been obtained from the probing cycle, we compare our mathematical eTTP prediction to such a simulation approach. We find that the simulation approach has an average error of 17.1%, which is 3.5 times higher than the error of the eTTP metric. We attribute this to the higher sensitivity of this approach to parameters such as *R*_0_, which cannot be estimated accurately from the probing cycle.

Summarizing, considering how the doubling time of resistant cells changed with the presence of a sensitive population allowed us to accurately predict which patients would benefit most from AT, and derive individualized estimates of TTP. Our mathematical metrics use data solely from the first treatment (probing) cycle, and outperform computational simulation approaches with the same data. We hope these metrics can serve as a mathematical biomarker to provide decision-support to clinicians and patients in deciding whether to receive adaptive or continuous therapy.

### 2.3 Personalizing Treatment

#### 2.3.1 Estimating the Personalized Treatment Threshold

For a potential AT patient, we can estimate the necessary parameters from the probing cycle and apply Eq. (4) to predict the optimal threshold size to follow in subsequent treatments, based on our chosen appointment interval (which is typically determined by clinical logistics/restrictions). This framework is visually summarized in Figure 2a. Applying this to the patients in our virtual cohort (Figure 4a), we observe significant heterogeneity in the optimal threshold between patients. For a given appointment interval, an optimal threshold for one patient will either be overly conservative or inherently risky for another.

**Figure 4:**
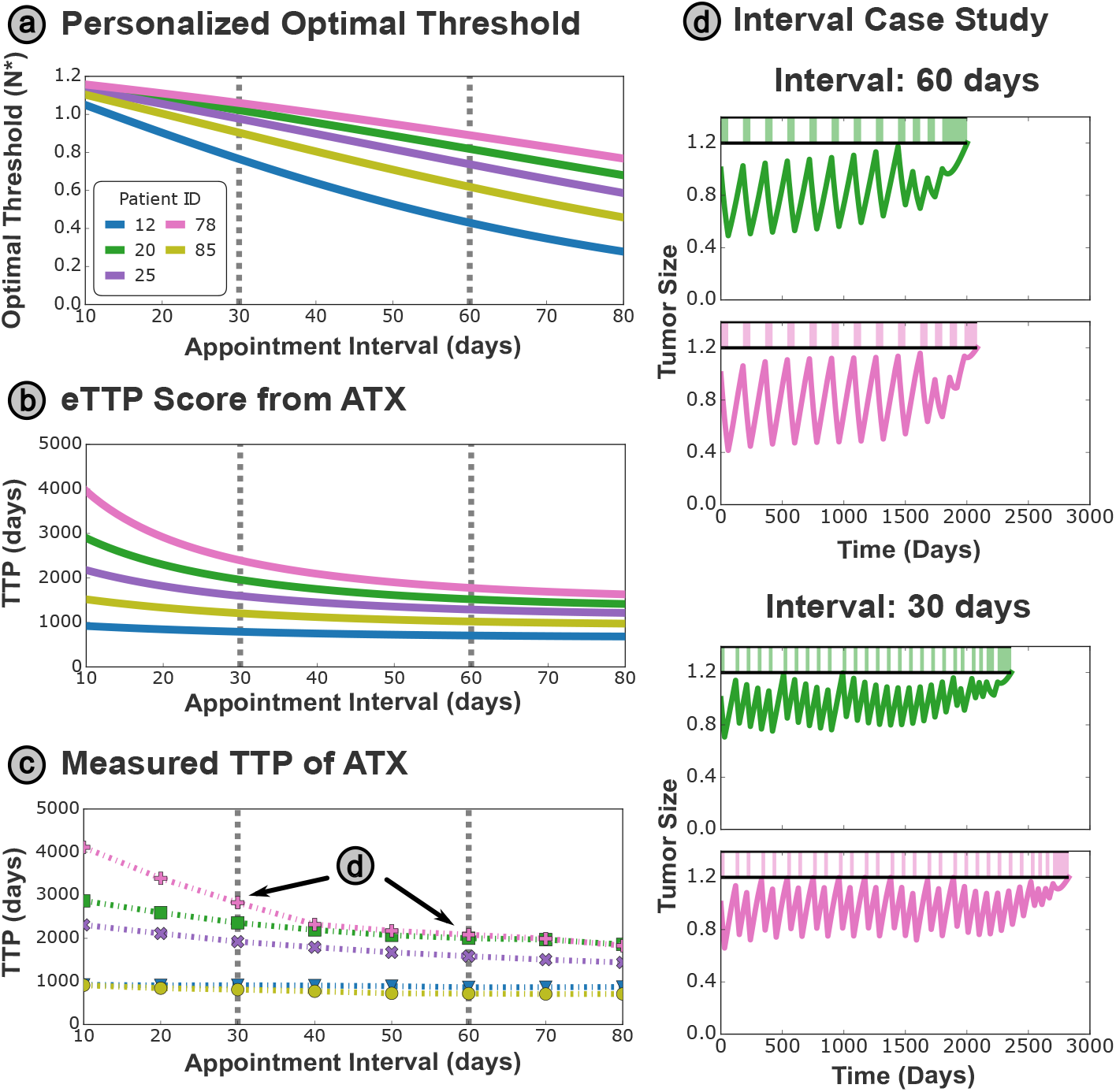
**(a)** Using parameters from the probing cycle fits, we can estimate the optimal threshold curves for each patient. **(b)** Combining the optimal threshold with estimates of both the doubling time and TTP under CT, we can compute the eTTP under the optimal strategy for each appointment spacing. **(c)** We find that the eTTP is highly predictive of the simulated TTP values from the virtual patient model, replicating both the ordering and approximate magnitude of the TTP benefit that each patient would expect. We see that some patients benefit significantly more than others, highlighting the importance of identifying these patients at the start of their treatment. **(d)** Highlighting case studies from monthly and bi-monthly intervals, we find that Patients 20 and 78 attain a similar TTP when treated bimonthly, but Patient 78 experiences a significantly greater benefit from monthly interventions.

In summary, patients exhibit significant heterogeneity in tumor dynamics highlights the pressing need for personalized adaptive therapy protocols. Based on our virtual tumor model (1), we analytically derive an optimal threshold that determines the best threshold size for AT based on the interval between clinical appointments, and show that this varies significantly between virtual patient profiles. The use of a personalized threshold mitigates the risk of early progression associated with previous approaches and accounts for patient heterogeneity to optimize TTP outcomes for each patient.

#### 2.3.2 Sensitivity to the Appointment Interval

Up to now, we have assumed that every patient is re-evaluated at the same interval (for example, many clinical guidelines foresee follow-up at standardized 2, 3 or 6 month intervals). However, we propose that in the context of AT, we should consider personalizing this follow-up frequency to maximize a patient’s benefit whilst subjecting them to a minimal required number of follow-up appointments. While we find that all patients benefit from a higher threshold size, and hence a shorter interval between appointments (see Figure 1c for an example), we also see that the magnitude of this benefit varies significantly between patients. It would therefore be clinically important to identify patients near the start of treatment that would benefit most from more frequent re-evaluation of their drug schedule. To do this, we apply the TTP prediction with the optimal threshold size determined solely by the appointment interval used for that patient.

In Figure 4b, we compute the eTTP for any patient undergoing AT at the optimal threshold for a given appointment interval. The qualitative trends in this plot (such as the ordering of both overall TTP and relative benefit from AT) strongly agree with the simulated outcomes for this test cohort, given in Figure 4c, confirming the accuracy of this predictive metric.

To exemplify the power of the eTTP metric, we may consider a typical bimonthly consultation (every 60 days), where in Figure 4d Patients 20 and 78 achieve approximately equal TTP. However, a significant disparity is observed when we halve the interval between treatments (to a monthly consultation); Patient 78 gains an additional 780 days until progression, while Patient 20 gains 350. This difference of over a year is stark, and while all patients do benefit from more frequent appointments, our approach may help clinicians prioritize patients (such as Patient 78) that will experience the greatest benefit.

### 2.4 Validation Dataset - Case Study

To conclude, we performed an independent validation of our methods on 3 patients from a clinical data set of castrate-resistant prostate cancer patients receiving adaptive androgen deprivation therapy (see Section 4.5 for details). Patient-specific tumor dynamics were predicted from the probing cycle (Figure 5a-b), and then benchmarked against a ‘ground-truth’ for each patient (which was derived from the patient’s full treatment history).

**Figure 5:**
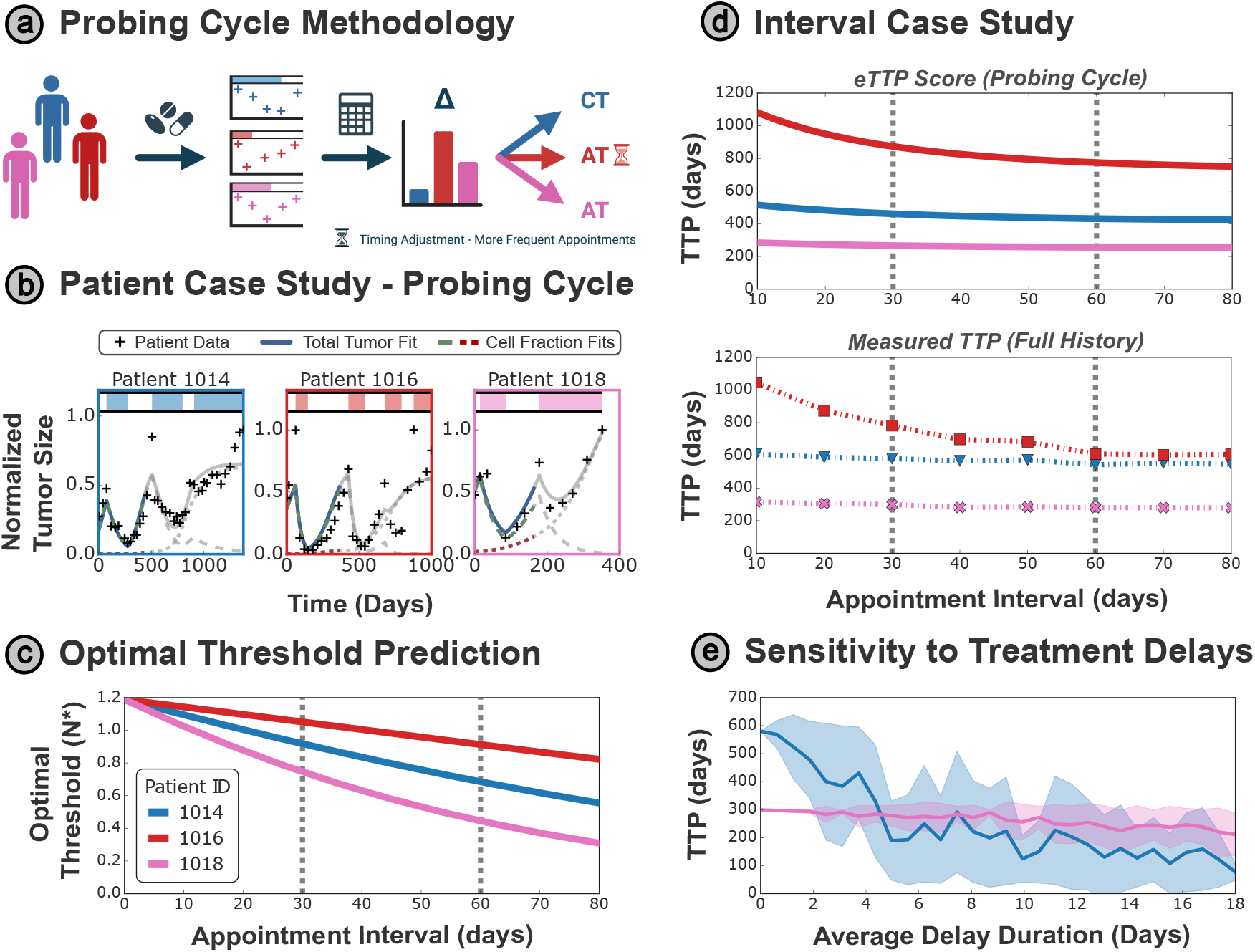
**(a)** New, unseen patients may undergo an initial probing cycle of AT50, to characterize their tumor dynamics, before being stratified into treatment protocols based on the Delta AT metric and eTTP score, extracted from the probing cycle data. **(b)** Data from Zhang et al. [18], unseen during methods development, is used to validate the approaches presented in this paper. The probing cycle fits are in color, with their continuation after the end of the probing cycle grayed out. **(c)** Using parameters from the probing cycle fits, we also see significant differences in the optimal strategy for each patient in this validation cohort. **(d)** The eTTP score (for AT treatment at the patient-specific optimal threshold for each appointment interval) identifies Patient 1016 as benefiting most from AT, which is borne out by simulated data, based on the entire patient history. This also highlights that Patient 1016 benefits most from more frequent treatment updates, as their TTP would increase the most under a transition from bimonthly to monthly appointments. **(e)** Considering the impact of unforeseen delays in treatment (modeled as a random delay sampled from a Poisson distribution), we plot the mean TTP for delays of a given average magnitude (the standard deviation of TTP under delays of a given mean duration is given by the shaded area). The separation between these curves motivates a further avenue for schedule personalization, with Patient 1014 being significantly more susceptible to these delays, and experiencing a much reduced TTP relative to Patient 1018.

Again, we observe significant heterogeneity in the optimal treatment strategy for each patient, with Patient 1016 benefiting from a significantly higher threshold size (*N* ^∗^) than Patient 1018 (Figure 5c). This is reflected in the TTP benefit expected for each patient under AT protocols with decreasing appointment intervals, where Patient 1016 also benefits the most (Figure 5d). This can be explained by the higher threshold size, and hence larger average sensitive cell population during treatment, that this patient is able to sustain - maximizing the competitive suppression of the resistant sub-population. This prediction is realized in the TTP outcomes based on the ‘ground-truth’ parameter set, confirming that Patient 1016 will indeed benefit the most from AT, and also from reduced appointment intervals to enable more effective competitive suppression. Indeed, while Patients 1014 and 1016 would expect a similar TTP under bimonthly appointments, Patient 1016 is seen to outperform Patient 1014 by over 200 days when this is upgraded to monthly appointments.

Finally, we consider another practical reality of real-world treatment: missed/delayed appointments. To explore the impact of unexpected delays in updating treatment plans - for example, due to medication shortage, illness, missed/re-scheduled appointments - we simulated such delays for the two patients with lower response to AT (Patients 1018 and 1014). To model scheduling disruptions, we sampled random delays from a Poisson distribution (note that this is strictly positive, so the patient cannot have an appointment earlier than expected) - for further details of this approach, see Appendix A.3. We find that Patient 1018’s TTP outcome is highly robust to such delays, with minimal decrease in the TTP even when the mean delay length is over half of the appointment interval (30 days in this case). By contrast, Patient 1014 is highly susceptible to such delays, with their TTP quickly halving and falling below that expected for Patient 1018 for delays of 5 days or more. In this case, there would be a strong case for additional monitoring of Patient 1018, and clinicians could decide to prioritize this patient if appointments are scarce. This demonstrates how mathematical modeling can identify additional risk factors associated with unexpected appointment delays to crucially support adaptive therapy decision-making in the clinic.

### 2.5 Wider Benefits of Adaptive Therapy

Throughout this paper, we have shown that personalized adaptive treatment schedules can improve TTP, and this benefit is significant for some patients. Moreover, we can predict which patients will benefit most from personalized AT, using parameters obtained from the initial treatment cycle, as well as identifying which patients will benefit most from more frequent consultations.

However, throughout this analysis, some patients have been left behind. While no one does worse under AT, patients such as 12 or 85 show minimum increases to their TTP in Figure 4c, which begs the question: ‘Is AT still worthwhile for these patients?’ While TTP is the most common metric for the success of adaptive treatment schedules [22, 29], it is not the only option. Treatment toxicity is a leading cause of failure to complete anti-cancer drug cycles [30, 31, 32], and around 20% of patients undergoing abiraterone hormone therapy for castrate-resistant prostate cancer either discontinue treatment or receive dose reductions [33]. Furthermore, the cost of therapeutic agents can place a prohibitive financial burden on many patients, providing an additional barrier to completing treatment schedules. For these reasons, it is also highly pertinent to consider the impact of AT schedules on the drug burden. Shown in Appendix A.4, we find that AT reduced drug burden significantly for all patients, providing a decrease in the average drug dose over treatment of between 20% and 60% for patients undergoing optimal AT with bimonthly appointments. This translates to lower accrued chemical and financial toxicity over the course of treatment, thereby increasing the completion rate of prescribed treatment courses in late-stage cancer; and, more importantly, providing a better quality of life for the patient.

## 3 Discussion

Recent clinical studies have demonstrated the potential of adaptive therapy to offer significant increases in TTP at a lower cumulative drug burden [18]. Given the number of ongoing adaptive therapy trials (in skin (NCT05651828 - BCC Trial), prostate (NCT05393791 - ANZADPT Trial), and ovarian (NCT05080556 - ACTOv trial) cancers) using a range of different AT protocols, it is of significant clinical interest to develop methods that can identify and personalize optimal treatment schedules. Here we focus on intermittent schedules (where we choose between administering and holding therapy) and look to derive optimal AT protocols personalized to individual patients.

All previous theoretical work (e.g. [20, 23]), has been based on continuous patient monitoring, whereas here we specifically focus on clinically realistic protocols. However, it becomes increasingly important to account for patient heterogeneity when the interval between tumor observations increases to clinically feasible timings, motivating the introduction of a personalized tumor size threshold for treatment decisions that adapt to variation in patient drug response. To ensure clinical applicability, where treatment schedules must be derived for patients with no treatment history on a given drug, we propose an initial probing cycle of the Zhang et al. [17] protocol (AT50), and then base all future personalization on data obtained from modeling this first treatment cycle (summed up in Figure 2a).

Using this approach, we propose two metrics - the Delta AT score and the eTTP - that can predict at the start of treatment the relative and absolute benefit of adaptive therapies respectively, helping inform treatment protocol selection and potentially motivating adaptive therapy referrals. During treatment, we are then able to determine the optimal threshold for each patient, based on both their tumor dynamics and clinical restrictions, such as the time interval between treatment consultations. Finally, we can also identify patients who would benefit from more frequent treatment re-evaluations, supporting efficient and evidence-based allocation of clinical time and resources. We also find that some patients have a greater susceptibility to unanticipated delays in treatment (Appendix A.3), and that this is dependent on the clinical appointment frequency, highlighting the need to incorporate clinical realities such as missed appointments that are typically neglected in idealized modeling scenarios.

This study specifically focuses on the well-studied Lotka –Volterra tumor model [21], however, we hope to extend this analysis to capture a wider range of model dynamics in future work. We also leave the quantification of uncertainty in our estimated tumor parameters from the first-cycle fits to future work, which we believe would support the robustness of these findings. Specifically, we acknowledge that patients in our clinical dataset only went through a small number of treatment cycles, and these approaches should be validated on patients with longer treatment histories, to ensure parameter values do not drift over time due to ongoing tumor evolution. However, we propose that these issues could be mitigated by continually refitting the models and recomputing metrics such as the optimal threshold size *N* ^∗^ as more data are accumulated, to ensure the accuracy and contemporaneity of the virtual patient representation, akin to the ADAPT protocol proposed by Strobl et al. [28].

In this paper, we assume that measurements of the tumor burden are only made at clinical appointments, when the treatment schedule is being re-evaluated. In reality, it is often possible for patients to undergo routine blood work at a local clinic, and this may allow for more frequent burden estimates than their appointments with a specialist oncologist. Particularly if additional blood tests are scheduled at the start of treatment, this would enable more accurate predictions of patient-specific tumor dynamics, and hence more accurate TTP predictions. Beyond this, it is possible to imagine future technological advances will allow for near-continual monitoring of blood metrics for tumor burden such as PSA, using home self-test kits similar to those commonly available to patients with diabetes. Such ongoing monitoring enhances the potential of ‘real-time virtual tumors’ which are continually updated based on the most recent data collected. These would provide clinicians with significantly greater certainty in optimal treatment predictions, and could reduce the need for blood tests in-clinic.

Through this study, we highlight the importance of accounting for clinical realities in mathematical derivations of optimal treatment schedules, and demonstrate that this can result in profound qualitative differences in the optimal treatment protocols obtained. Here, the inclusion of discrete patient monitoring demonstrates that prior standardized treatment schedules do not adapt to patient heterogeneity, motivating the need for personalizing adaptive treatment strategies. We further illustrate the power of the probing first-cycle model fits to capture heterogeneity in patient dynamics, enabling treatment personalization for new patients and leading to safer and more effective treatment schedules. Overall, we hope this work has highlighted the importance of personalized treatment schedules that explicitly account for patient heterogeneity, and the power of mathematical models to capture, analyze and facilitate this personalization.

## 4 Methods

### 4.1 Lotka– Volterra Model

Growth and drug-response dynamics of a partially drug-resistant tumor were represented by the two-population Lotka–Volterra model [34, 35] introduced by Strobl et al. [21], where variation in drug sensitivity is represented through separate drug-sensitive and -resistant cells, denoted by *S*(*t*) and *R*(*t*) respectively for time *t*:

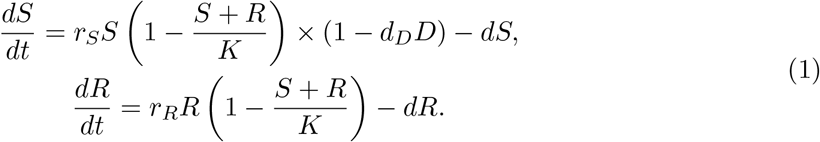

Both species follow a modified logistic growth model with growth rates *r*_*S*_ and *r*_*R*_ and a common natural death rate *d*, where the total population (rather than the species population) is modified by the shared carrying capacity *K*. As drug-resistant cells typically have a resistance cost [36], we consider the regime where *r*_*S*_ ≥ *r*_*R*_, including the case where there is no such cost. Treatment is assumed to kill sensitive cells at a rate that is proportional to the population’s growth rate (the Norton–Simon model [37]) and the drug concentration, *D*(*t*) (with proportionality factor *d*_*D*_). Note that this model does not account for resistance acquisition during treatment; Viossant and Noble [20] demonstrated that the inclusion of genetic mutation in simple models does not significantly alter the response to therapy.

Parameter values were adopted from Strobl et al. [21] and are given in Table 1. Note that throughout this paper, we will use *N* to denote the total tumor size - i.e. *N* = *S* + *R*.

### 4.2 Adaptive Therapy

We benchmark all adaptive treatment schedules against the following two standard protocols:

1. **CT** – Continuous Therapy (Standard of Care):

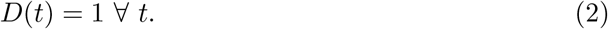
2. **ATX** – Treatment is given until a decrease to X% of the initial size (*N*_0_) is achieved, then withdrawn until the tumor returns to its initial size:

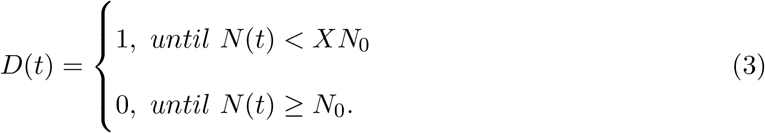

A value of *X* = 50 % was used in the pilot AT clinical trial (AT50) by Zhang et al. [17, 18]. In all treatment schedules, a drug dosage of 1 corresponds to the maximum tolerated dose. Treatment outcomes from these schedules are compared according to their TTP, where progression is defined as a 20 % growth from the initial size (i.e. 1.2*N*_0_).

### 4.3 Calculating the Optimal Threshold Size

The optimal appointment interval for AT50 (as motivated in Section 2.3.1) is directly determined by the growth characteristics of the tumor - specifically, failure occurs when the tumor can grow from the treatment threshold size to the progression threshold size in the time interval *τ* between consecutive appointments. The authors have separately explored this phenomenon [24], where they derive the following expression for the optimal treatment threshold *N* ^∗^(*τ*) for a given appointment spacing (*τ*):

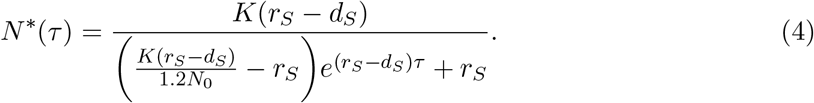

### 4.4 Prediction Metrics

#### 4.4.1 Delta AT Score

The natural timescale for the growth of the resistant sub-population may be captured by the doubling time of resistant cells (from an initial population of size *R*_0_). In Appendix A.2.2, we derive the following expression for the doubling time, in the presence of a fixed population of *κK* sensitive cells (i.e. some fraction *κ* of the carrying capacity *K*, where *κ* ≤ 1):

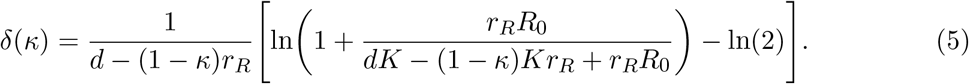

We therefore write the benefit (in doubling time) derived from the presence of *κK* sensitive cells (hereafter referred to as the Delta AT score) as

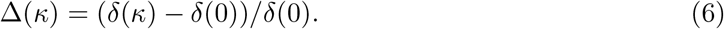

This is evaluated at a fixed fraction of the carrying capacity for all patients - *κ* = 0.2 is used in this paper.

#### 4.4.2 eTTP Score

To generate the estimated TTP (eTTP), we first simplify the mathematical tumor model by assuming the sensitive population is maintained at a fixed level over the AT protocol. This provides a fixed level of competitive suppression on the growth rate of resistant cells, which we can compute on a patient-specific basis. Comparing this to the TTP expected when there are no sensitive cells present to halt the growth of the resistant cells, we can therefore compute an estimate for the overall TTP. To ensure this prediction is accurate within the clinical context of fixed intervals between appointments (as opposed to continuous monitoring of the tumor), which allows the tumor size to overshoot the designated threshold sizes between appointments, we add an additional correction term (*ζ*) to our estimate of the effective sensitive population size 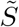 during treatment. A visual summary of this derivation is given in Figure 3a - the full derivation is provided in Appendix A.2, and results in a final expression for the eTTP of:

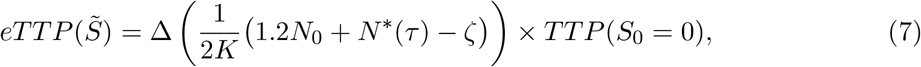

where the overshooting term *ζ* is given by:

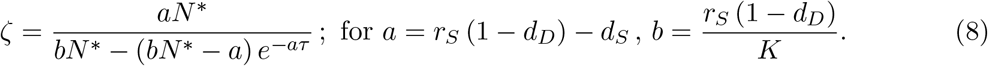

### 4.5 Patient Data

Virtual patient profiles were based on two publicly-available data sets of longitudinal PSA records for patients with prostate cancer undergoing androgen deprivation therapy. Data from Bruchovsky et al. [25] were used in Sections 2.2-2.3, while independent data from Zhang et al. [18] were used as validation in Section 2.4.

Enrollment into the Bruchovsky trial required histologically-confirmed adenocarcinoma with a rising serum prostate-specific antigen (PSA) level after radiotherapy and no evidence of distant metastasis. Patients were subject to an intermittent therapy regime, where treatment in each cycle consisted of cyproterone acetate given as lead-in therapy for 4 weeks, followed by a combination of leuprolide acetate and cyproterone acetate, up to a total of 36 weeks. Serum PSA and testosterone levels were monitored every 4 weeks, and fixed PSA thresholds of 4*µ*g/L and 10*µ*g/L were used to determine treatment cycles.

We generated virtual tumor profiles by fitting Equation (1) to each patient’s longitudinal data, minimizing the root mean squared difference between the normalized PSA measurements, and simulated tumor volumes. Full details of this fitting process are given in Appendix A.1.1. From this, we formed an example test cohort composed of 5 patients who progressed clinically and simulated the application of conventional and adaptive treatment strategies to these patients.

The validation dataset used in Section 2.4 is taken from a recent clinical trial of adaptive therapy in castrate-resistant prostate cancer [18]. Patients in the study cohort were subject to conventional AT50, wherein abiraterone treatment was stopped when PSA was less than 50% of pre-treatment value and resumed when PSA returned to baseline. Patients could be enrolled in the study after achieving a greater than 50% decline from their pre-abiraterone PSA levels. To generate virtual patient models from each patient’s longitudinal PSA records, the same fitting procedure was applied as above.

## A Appendices

### A.1 Fitting Procedures

Motivated by the fitting approach taken by Strobl et al. [21], all fitting was conducted by minimizing the root mean squared difference between the prostate-specific antigen (PSA) measurements and the predicted tumor size, both normalized to the initial size. This was applied using the Levenberg–Marquardt algorithm, implemented using the lmfit package in python [42]. Patient profiles were excluded if no satisfactory fit could be obtained (*r*^2^ *<* 50% or failure to capture later stage cycles), typically due to insufficient PSA measurements, insufficient measurement frequency or no evidence of relapse.

#### A1.1 Virtual Patient Model

To generate a virtual patient model for each patient that can act as a ground truth to validate the predicted behavior of our mathematical biomarker, we fit to the full treatment history of each patient. While these patients were not subject to a true AT50 protocol, we will use the data from the first complete treatment cycle as a de-facto probing cycle for future analysis.

For each individual, we fit the parameters: *r*_*S*_, *r*_*R*_, *d*_*S*_, *K*, along with the initial tumor cell density *N*_0_ and resistant cell density *R*_0_. While this fitting is done in a single step, the identifiability of the model parameters relies on an inherent timescale separation of this models’ dynamics. The initial treatment response is predominantly determined by the parameters for the sensitive population, while the fraction of resistant cells is negligible, whereas the late time growth rate and TTP is controlled by the resistant cell parameters.

While these patients were not subject to a true AT50 protocol, we will use the data from the first complete treatment cycle as a de-facto probing cycle for future analysis. Given the limited information available in the probing cycle, the fitting procedure for the probing cycle data differed substantially from that used for the virtual patient model, and is outlined in Appendix A.1.2.

#### A.1.2 First Cycle Parameters

When predicting patient-specific tumor dynamics from PSA readings in the first treatment cycle only, we do not have access to these two timescales. Given the extremely limited information available in this scenario, the model is no longer fully-identifiable, and we are unable to extract all the parameters detailed in Section A.1.1. For this reason, the parameters *r*_*S*_, *r*_*R*_, *R*_0_ are set to values from the literature quoted in Table 1 (or at the lower end of ranges given), chosen due to the reduced variance of these parameters across the virtual patient cohort. Should prior clinical information (unrelated to the current patient cohort under treatment) be available, these population-wide values could instead be determined by average parameter values from historical cohorts.

Furthermore, to ensure the identifiability of the model in this reduced information context, we also exclude the initial tumor density *N*_0_ from the parameter fitting; instead this is set equal to the first recorded PSA measurement (normalized relative to the maximal PSA value recorded). We therefore only fit for the parameters *d*_*S*_ and *K*, which are identifiable from the first treatment cycle (comprised of both the response to the initial treatment period, and subsequent off-treatment rebound back to the original tumor size).

### A.1 TTP Prediction

Evaluating the TTP analytically would require solving a coupled, nonlinear ODE system with a piecewise drug term, which is not analytically tractable. Instead we can derive an approximation by first considering the simpler case of a fully resistant tumor, which we can solve analytically in Section A.2.1 to obtain a base-line (worst case) TTP in the case of zero sensitive suppression. Repeating this derivation for a fixed, non-zero sensitive population, we then compute in Section A.2.2 the benefit conferred by the presence of sensitive cells. These suppress the growth of the resistant cells, and we introduce a new metric (the Delta AT metric) to quantify the impact on the resistant cell growth dynamics.

Given the TTP is dependent on the size of the sensitive population over the course of treatment, we introduce a function *TTP* 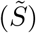 to denote the TTP expected in the presence of a fixed sensitive population of size 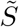. We can use the relative increase in the growth timescale of the resistant population to scale *TTP* (0) to an overall prediction of the TTP (*T* (*κ*)), based on the assumption that there is a sensitive population of average size *κK* present during treatment. Finally, in Section A.2.3 we compute this average tumor size, assuming that the population will be constrained between the progression limit (1.2*N*_0_) and the chosen treatment threshold *N* ^∗^. For discrete time intervals between appointments, the tumor size is not perfectly constrained between these limits - the tumor size will overshoot the lower threshold and continue to decrease until a treatment holiday is applied at the next appointment time. Similarly, to avoid the early tumor progression, the threshold tumor size is designed to ensure treatment stops short of the progression limit, instead of overshooting it within the next appointment interval. We therefore also quantify this overshooting difference, and correct the average size for this.

This approximation is based on various assumptions, primarily that the average competitive suppression generated by a varying sensitive cell population is approximately equal to the competitive suppression generated by the average size of the sensitive population. While we motivate the reasoning behind these assumptions, this is intended as an approximation of a quantity that is otherwise analytically intractable, and not a formal derivation of this quantity in some limit. This approach is instead validated through simulations in Figure A.2.4, where it is shown to be a good proxy for simulated TTP across the virtual patient cohort.

#### A.2.1 Resistant-Only TTP

First we consider the non-coupled ODE for a fully resistant tumor of initial size *R*_0_:

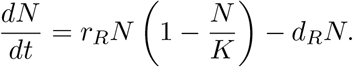

Because the tumor is fully resistant, this expression holds whether or not drug is present. Evaluating this separable differential equation, we obtain:

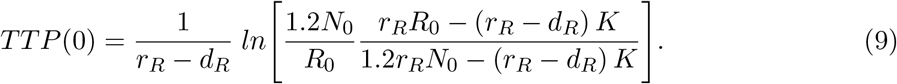

Note that we write the progression limit as 1.2*N*_0_ instead of 1.2*R*_0_ - while *R*_0_ ≡ *N*_0_ in this case, we will later apply this result to tumors with both sensitive and resistant cells, where this equivalence does not apply.

#### A.2.2 Delta AT Metric

We can consider the impact of sensitive cells on the TTP mathematically, by contrasting a zero suppression case (where *S*(*t*) = 0 ∀ *t*) with a non-zero suppression case (where *S*(*t*) = *κ* ∀ *t*, with 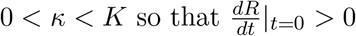.

To characterize the growth rate of sensitive cells, let us consider the doubling time (*δ*(*κ*)) of the resistant sub-population in the presence of *κK* sensitive cells. We may integrate Equation (1) with fixed *S* = *κK* between the original tumor size *R*_0_ and some final tumor size *R*_*f*_ to obtain the expression (duplicated from (5)):

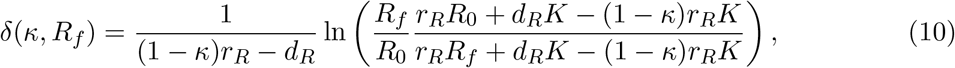

which, with *R*_*f*_ = 1.2*N*_0_, is equivalent to (9) in the special case of *κ* = 0.

However it is more intuitive to consider the doubling time of resistant cells, by setting *R*_*f*_ = 2*R*_0_. We use this to generate the predictive Delta score generated in Section 2.2.1, thereby removing any assumptions about the overall tumor size or progression limits. In this case, (10) reduces to (duplicating (5):

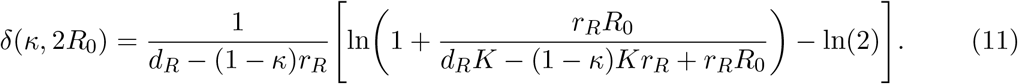

Finally, we summarize the relative benefit conferred by this competition suppression through Δ(*κ*) - i.e. the proportional increase in the growth time from *R*_0_ to *R*_*f*_ due to the presence of *κK* sensitive cells, which we call the Delta Score (replicating (6)):

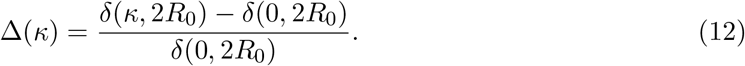

From Eq (5), we may also identify patient characteristics that contribute to an increased Delta AT metric, and hence an increased benefit from AT (Figure 6a). Naturally, an increased cost of resistance (which is defined by 1 − *r*_*R*_*/r*_*S*_) corresponds to enhanced suppression of the resistant population as the sensitive population has a greater relative fitness advantage off-treatment, increasing the benefit of AT. This impact is more significant when the resistant fraction is larger (or the carrying capacity is smaller), as the tumor is closer to the carrying capacity of the system, and hence more susceptible to competitive pressure. Finally, increased cellular turnover (defined by *d*_*S*_*/r*_*S*_) has a similar impact, also increasing the resistance fraction and hence the inter-species competition. Note that (9) does not depend on the initial tumor size (only the initial resistant population), and hence this is not included in the sensitivity analysis. Similarly, given that the doubling time is derived in the case of perfect resistance, parameters associated with the tumor’s drug response such as *d*_*D*_ are also not considered.

**Figure 6:**
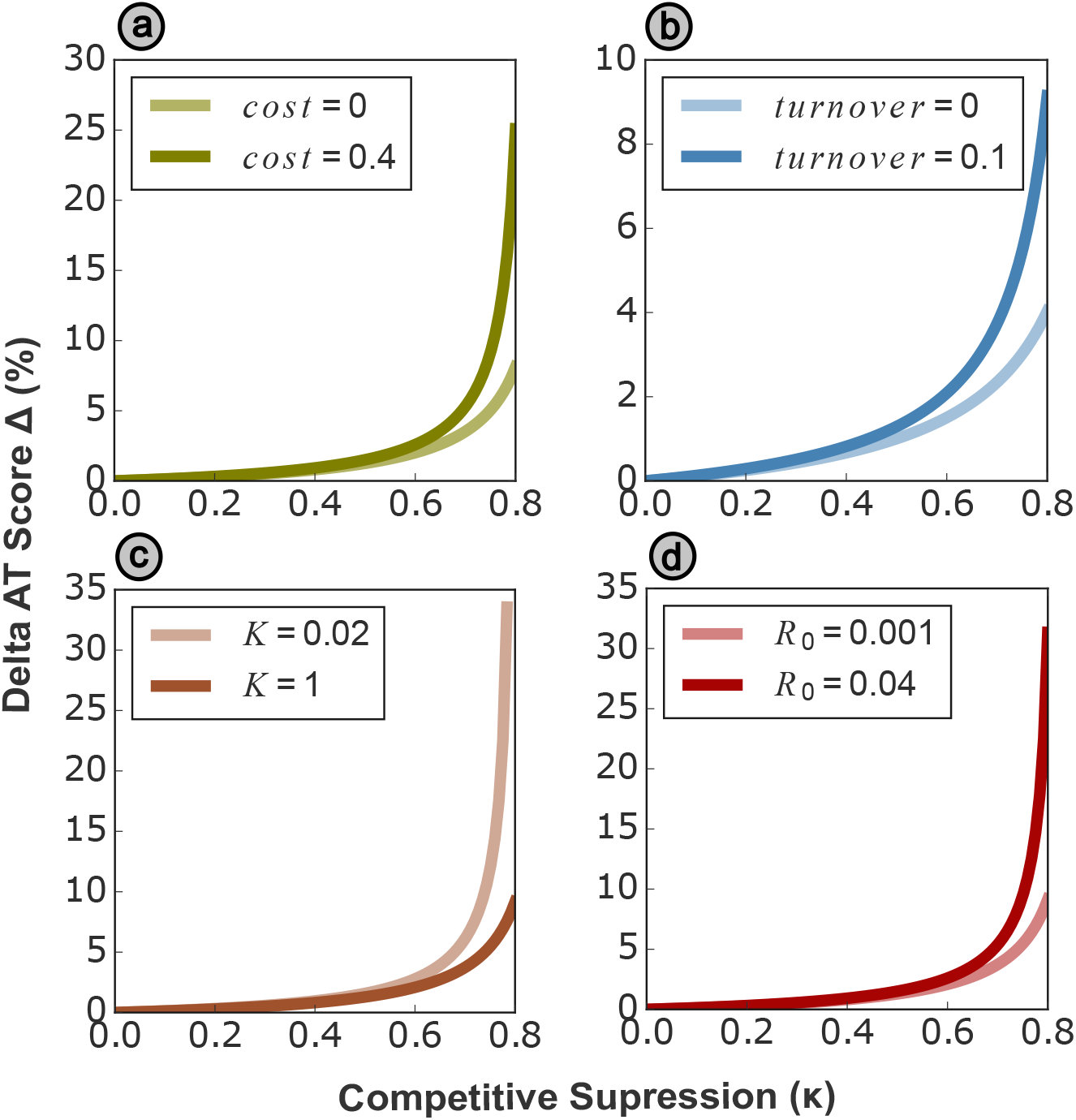
Conducting a basic local sensitivity analysis of the Delta AT metric Δ(*κ*) (5), we can demonstrate its dependence on the four key model parameters: the cost of resistance, the cellular turnover, the carrying capacity and the initial resistance fraction. We can predict the benefit of constant sensitive population sizes for differing values of *κ* (the size of the constant sensitive population), with (*cost, turnover, K, R*_0_) = (0.1, 0.1, 1, 0.001) unless otherwise stated. We find that increased cost and turnover enhance the benefit of a sensitive cell population (as they increase competitive suppression of resistant cells), while both an increased initial resistant fraction and a reduced carrying capacity also increase the benefit (because the cells are closer to carrying capacity and so more affected by inter-species competition).

#### A.2.3 Average Sensitive Population & Overshooting Correction

Finally, scaling the baseline TTP by the Delta AT score, we may estimate the TTP (*T* (*κ*)) as:

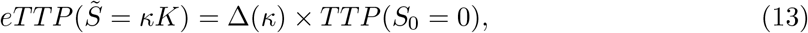

where 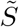 denotes the average size of the sensitive cell population over the entire treatment history. Because sensitive cell dynamics vary with the presence of drug, unlike the resistant cells used to derive *TTP* (*S*_0_ = 0), the value of 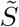 (and hence overall TTP prediction) is dependent on the treatment protocol considered. To approximate 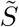 in ATX schedules, we take the arithmetic mean of the progression limit (1.2*N*_0_) and threshold threshold (*N* ^∗^).

However this does not account for overshooting these threshold values due to the discrete interval *τ* between appointments - an example of this phenomena is shown in Figure 3. When there is a single threshold value, this effect is negligible (as overshooting above and below the threshold are assumed to cancel out over sufficiently many treatment cycles). However, in conventional ATX where we can consider the tumor size to be trapped between the lower threshold size, and the progression limit, the tumor size will overshoot below the threshold tumor size, while never reaching the progression threshold. We account for this effect by computing the average overshooting of the sensitive population past the tumor size threshold (*N* ^∗^) during the time interval between appointments. To compute this, consider the following simplified equation for the tumor under constant drug treatment, taken from the Lotka – Volterra system (1) under the assumption of a negligible resistant population:

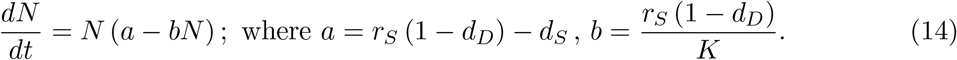

Integrating this over the full appointment interval *τ* (i.e. considering the worst-case scenario that the tumor passes the threshold size immediately after the most recent appointment) gives the maximum possible overshoot (*ζ*) of the tumor size *N* (*t*) past the treatment threshold *N* ^∗^ in time *τ* :

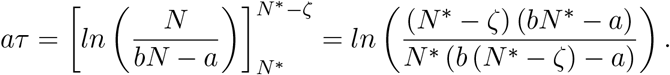

We can rearrange this to obtain an explicit form for the maximal overshoot:

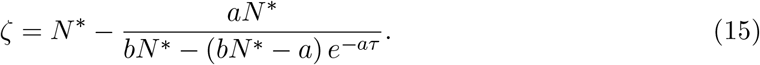

We estimate the average tumor size by subtracting the average correction term *ζ/*2 from 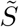 (both scaled by the carrying capacity *K*), to give the final TTP prediction as (replicated from (7)):

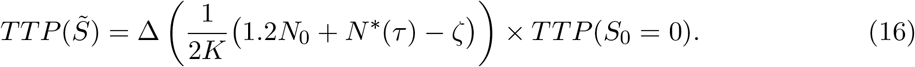

Note that this correction is only required for AT protocols based on a treatment window (such as ATX) implemented with discrete monitoring (i.e. a non-zero appointment interval *τ*). When a single threshold is used (such as the personalized treatment threshold *N* ^∗^), we assume that overshooting above and below the threshold cancel out over sufficiently many cycles. To approximate the average size of the tumor as the optimal threshold size *N* ^∗^, we also implicitly assume the duration of the final treatment cycle (wherein the primarily-resistant tumor grows from *N* ^∗^ to the progression limit 1.2*N*_0_) is short relative to the overall duration of the AT protocol. Based on this, we can simplify (16) to:

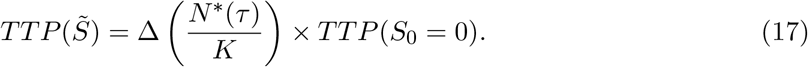

#### A.2.4 Validation of eTTP

To validate these approaches, we compare the eTTP to the simulated TTP under a range of treatment protocols. We use profiles from the virtual patient cohort, and make predictions using the same parameter set as we use for simulation.

We first consider a CT protocol, wherein sensitive cells are quickly eliminated by continuous drug treatment, and the tumor behaves as the fully-resistant tumor considered in Section A.2.1. Based on (9), we obtain accurate estimates of the TTP with a mean absolute error of 3.4%, though this is due to a consistent under-estimation of the true TTP. This error may be fully attributed to the competitive suppression by the sensitive population at early times before it is eliminated. While this is difficult to account for analytically, we discuss and quantify this error further in Section A.2.5. However, the highly accurate estimation of the TTP we obtain in Figure 7 demonstrates the validity of this approach, despite our neglect of the impact of sensitive cells on this baseline TTP.

**Figure 7:**
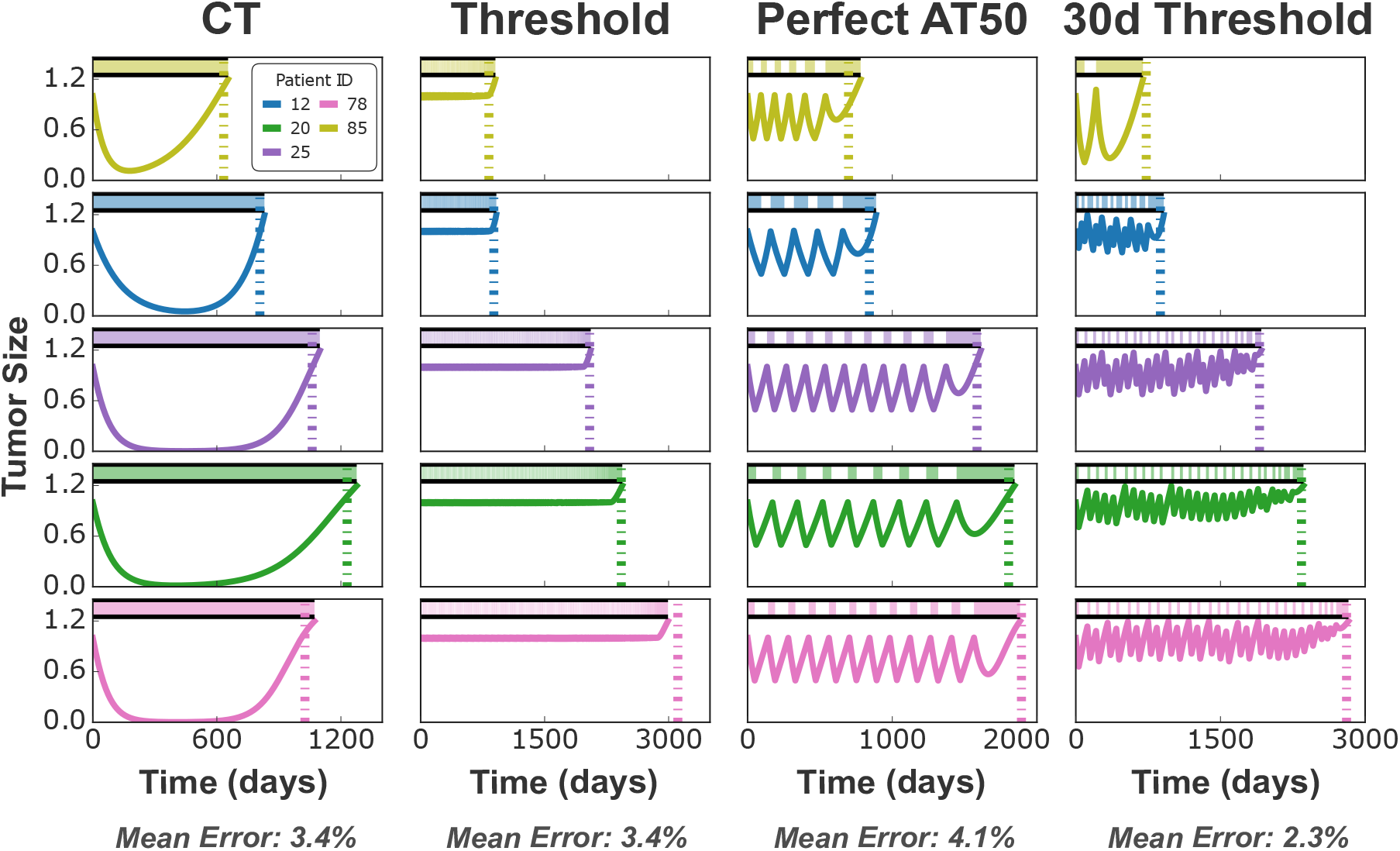
We validate the eTTP scores (dashed lines) against four simulated treatment protocols (solid lines) of increasing complexity. We first analyze a CT protocol, which may be approximated in a similar way to the resistant-only TTP. Following the approach outlined under the Delta AT metric derivation (Section A.2.2) we then consider to a threshold-based schedule, akin to a fixed sensitive population. We then extend this to perfect AT (with continuous monitoring to prevent overshooting), where the average sensitive population 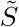 is given by the average of the threshold sizes. Finally we may account for overshooting by considering a clinically-realistic threshold schedule, where treatment is only re-evaluated on 30 day intervals. In all cases, the eTTP approach is able to predict the TTP with less than an average error of 5% across all patients in the virtual cohort.

We then introduce the Delta AT metric, which proportionally scales the baseline TTP according to the average sensitive population under treatment, to give the overall TTP prediction (the eTTP). This approach is most easily validated using a threshold protocol, wherein treatment is modulated to obtain a constant overall tumor size. In Figure 7, the threshold protocol holds the tumor at the original tumor size, until progression is driven by growth of the resistant population. The final resistant population is typically much larger than the initial resistant population (*R*_*f*_ ≫ *R*_0_), and hence may be on the order of the carrying capacity of the tumor *K*, changing the growth dynamics of the tumor. Evaluating the Delta AT metric at *R*_*f*_ = 2*R*_0_, as used in Section 2.2.1, is therefore not appropriate when trying to obtain an absolute prediction of the TTP.

Under the CT treatment protocol, the sensitive population is no longer eliminated, and hence the final tumor is comprised of a mix of sensitive and resistant cells. In simulations we observe that sensitive and resistant cells are present in approximately equal proportions, and hence the resistant cells have a final population size of 0.6*N*_0_ (half the total tumor size) at progression. While resistant cells typically exist in slightly higher proportions than sensitive cells in these simulations (see Figure 1a for an example of this), we find that this bias is counter-balanced by a reduction in the competitive suppression driven by the sensitive cells when they are depleted at late times. In further analysis for the eTTP score, we therefore use *R*_*f*_ = 0.6*N*_0_ unless otherwise stated.

While the fixed total tumor size in the threshold protocol allows us to approximate the average sensitive population 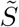 as approximately equal to the threshold size, 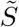 is harder to compute for range-based implementations of AT such as ATX. This protocol bounds the tumor size between two limiting sizes, and we compute 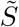 as the arithmetic mean of these limits. While this is only an accurate time-average of the tumor size in the simplistic case of linear tumor growth dynamics, we assume that the mean acts as a good approximation of 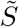 for the tumor dynamics simulated here (for example the oscillations in tumor size under the ATX schemes of Figure 1a are approximately linear). To validate this assumption, we consider a ‘perfect’ implementation of AT50, where tumor size is monitored continuously and does not overshoot the limits prescribed in (3). Again, we find that the eTTP prediction correctly orders the simulated TTP between all patients in the virtual cohort, and predicts the absolute value of the TTP with less than a 5% mean absolute error (Fig 7).

However, practical implementations of AT cannot monitor the tumor size constantly, and therefore the tumor size will inevitably overshoot the threshold sizes between appointments. This means that a simple arithmetic mean of the limit sizes may no longer be a good approximation of 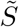, and so we account for this overshooting in an additional term in Section A.2.3. To exemplify the application of this correction term, we consider a threshold-based treatment protocol, with a 30 day appointment interval (i.e. treatment may only be re-evaluated at 30 days intervals). In Figure 7, we see this inconsistency in tumor size between oscillations, over-shooting the threshold size as a result of the appointment interval. However the eTTP accounts for this effect to produce highly accurate TTP predictions with an average error of 2.3%.

#### A2.5 Role of Sensitive Population in CT

In Section A.2.1 we assumed that the effect of the sensitive population is negligible when predicting the TTP under CT. Specifically, given that a non-zero sensitive population suppresses the growth of the resistant population, we have assumed that the sensitive population exists for negligible time before it is eliminated (or at least reduced to a negligible size). However this is not true — the sensitive population persists for a finite time and the suppression that it exerts during this period is responsible for the prediction inaccuracy observed under the CT protocol in Figure 7.

We considered two separate simulation approaches to confirm this was the source of the observed discrepancy, first artificially neglecting competition by setting the carrying capacity *K* to a sufficiently large value (*K* = 100 was found to suffice). As this also modifies the growth dynamics of resistant cells (changing the growth characteristic from logistic to exponential), we considered a second validation approach where the sensitive population was eliminated all together (i.e. *S*(*t* = 0) = 0). In both cases that we found that the eTTP metric was able to predict the exact TTP to the nearest day, confirming that this competitive suppression exerted by the sensitive cells was the source of the small observed discrepancy in the CT panel of Figure 7 (results not shown).

We may consider a timescale separation approach to quantify the impact of competitive suppression more formally. For small initial resistant fractions *R*_0_, we find that the sensitive population may be effectively eliminated (i.e. *N* (*t*) *< N*_0_*/*100) under CT before the resistant fraction reaches a non-negligible size. Such dynamics are seen in 4/5 patients in Figure 1b, where the sensitive population may be reduced to undetectable levels before the resistant cells reach a measurable cell fraction. For this reason, we consider separating the treatment response into two distinct timescales: a ‘fast’ timescale where the sensitive population is depleted, and a ‘slow’ timescale where the resistant population grows to replace it, without feeling the effects of the (now-extinct) sensitive population.

Crucially, on the fast timescale, the resistant population size is negligible and so does not exert any competitive pressure on the sensitive population. However the resistant population is suppressed by the much larger sensitive population, and this suppression during the fast timescale at the start of treatment results in the small delay to simulated progression that we observe under the CT protocol of Figure 7, relative to our prediction that does not account for this effect.

First considering the fast timescale, we may solve the sensitive population dynamics analytically, as they are effectively independent of resistant cells. We have already considered this system when considering the full tumor with negligible resistance in (14), which we repeat below in terms of *S* only for clarity:

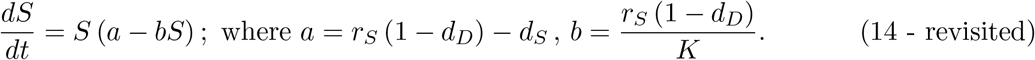

However, for the sake of later analytic tractability, we will assume that the effect of the drug kill on sensitive cells is much greater than the effect of competitive suppression while the drug is active (i.e. that *a* ≫ *bS* as *a <* 0) — quite a reasonable assumption in this system given *d*_*D*_ *>* 1 and hence *b <* 0. This gives the simplified equation *dS/dt* = *aS*, which has the solution *S*(*t*) = *S*_0_*e*^*at*^, where *S*_0_ := *S*(*t* = 0) (i.e. the initial sensitive population).

We may then substitute this expression into the equation for the resistant cell dynamics to consider the impact the non-zero sensitive population has on the fast-timescale growth dynamics of the resistant population:

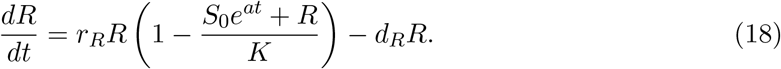

During the fast timescale (at small values of time *t*), *S*_0_*e*^*at*^ ≫ *R*(*t*) for sufficiently large *S*_0_*/R*_0_, and so we neglect the competitive pressure that resistant cells exert on themselves. Note that this approximation only applies on the fast timescale, and our original calculations in Section A.2.3 account for the logistic aspect of the resistant cell growth dynamics on the slow timescale, when *R*(*t*) ~ *N*_0_. We rewrite (18) as:

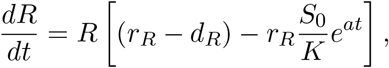

which we may solve to give:

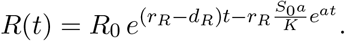

However, it is not possible to rearrange this expression for a closed form expression for *t* comprised solely of elementary functions. We must introduce the Lambert function *W* (also known as the product logarithm) to write *t* as:

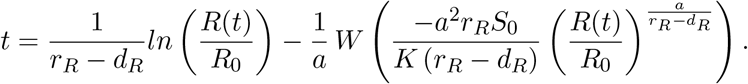

To compute the discrepancy *t*^′^ in TTP as a result of competition, we may recall that the corresponding expression for *t* in the competition-free limit (i.e. where *S*_0_*/K* → 0, such that 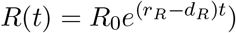 is:

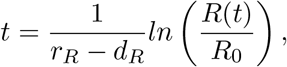

and hence we may write *t*^′^ as:

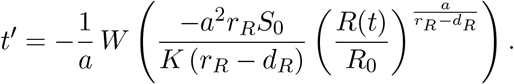

We may evaluate this numerically, using the scipy implementation of Corless et al. [43]. Considering the dynamics of the resistant population over the fast timescale, we evaluated *t*^′^ for *R*(*t*) = 2*R*_0_, and found that this correction accounted for less than a day for all patients (in comparison to doubling times of approximately two months), and corresponded to an error less than 0.5%. This relative error is reduced over longer timescales, and therefore can be considered insignificant in the overall eTTP prediction. While this approximations assumes perfect timescale separation, the magnitude of the discrepancy in Figure 7 shows that it is still relatively small (compared to the uncertainty in estimating probing cycle parameters when practically applying the eTTP metric) in simulated treatment response, and therefore does not impair the implementation of this metric.

### A.3 Robustness to Variation in Appointment Interval

In Section 2.4, we consider the robustness of different AT protocols to unexpected increases in the interval between appointments (be that due to clinical scheduling or unforeseen patient circumstances). We simulate the TTP under an AT protocol, where each treatment/holiday period is extended by a random delay, sampled from an exponential distribution. This delay is resampled for each appointment, and is independent of whether the patient is currently under treatment or under a treatment holiday. The mean of the delay distribution is given by the baseline interval length scaled by the noise factor — this delay is plotted for different noise factors in Figure 8. Simulations vary from the case with zero noise to a maximal-noise situation, where the mean of the delay distribution is given by half of the baseline appointment intervals (15 days for monthly appointments). We consider the impact of delays on both monthly and bimonthly AT protocols for each patient, where the threshold size used in each case is optimized for both the patient and appointment interval. Given the stochastic nature of the delay sampling, we average over 20 evaluations of the AT protocol for each noise fraction, and plot the standard deviation of these as the shaded area.

**Figure 8:**
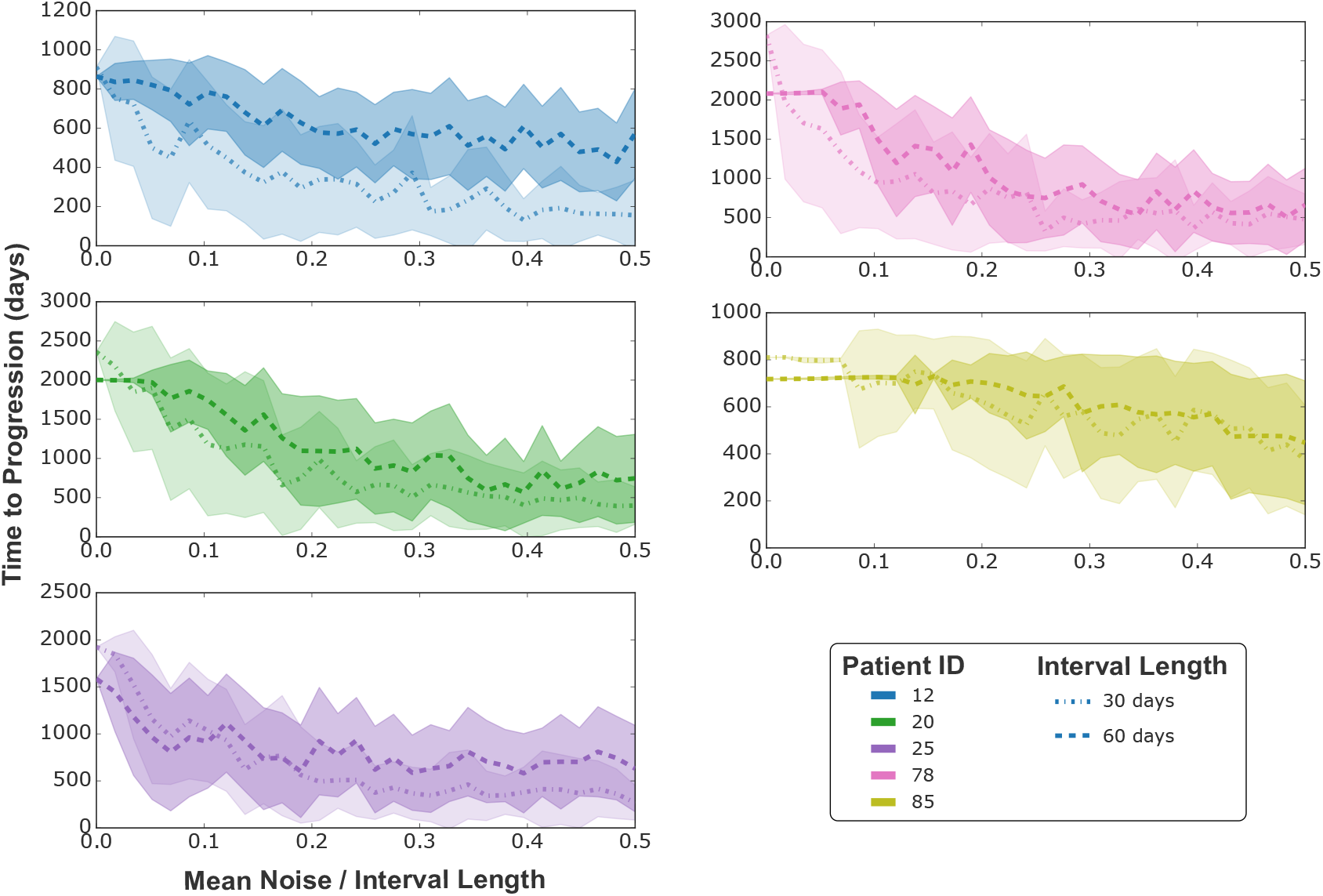
The average TTP under the analytically optimal strategy, when treatment decisions are subject to an unanticipated random delay. Delays are sampled from an exponential distribution (with the mean of the noise varied along the x axis, normalized by the interval length), and the mean and standard deviation of the TTP is computed across 20 iterations. We find that some patients have much greater reductions in their TTP for decision delays of a given magnitude than others (with Patient 85 being particularly robust to unexpected delays in treatment), and that this sensitivity to delays is also dependent on the appointment interval, motivating the need for clinicians to consider this personalized risk profile when scheduling follow up appointments, and allocating limited clinical resources.

As expected, all patient’s TTP outcomes decrease as the average magnitude of the delay increases, however the extent of this decrease varies markedly between patients: Patient 25 experiences significantly reduced TTP with delays less than 10% of the interval length, whereas Patient 85 can tolerate significantly longer delays without a reduction in TTP. The baseline appointment interval of the protocol matters too - we find that Patient 12 is more robust to delays while under an AT protocol with a baseline interval of 60 days, and Patient 78 actually has a longer TTP under a 60 day interval for small delays.

### A.4 Reduction in Combined Drug Use

In Section 2.5, we emphasize that TTP extension is not the only relevant metric for the success of AT protocols, and also consider the reduction in drug required. This is quoted as an average dose over the course of treatment (compared to unity for continuous therapy), as patients who also benefit from an enhanced TTP will have this drug spread over a longer time period. Note that the actual daily drug dose remains binary in this clinical context - an average daily drug dose of 0.5 means that the patient will be prescribed the drug for approximately half the time they are under the AT protocol.

First considering so-called ‘Perfect’ AT, where tumor monitoring is continuous and changes to the treatment schedule are instantaneous, we see in Figure 9a that all patients benefit from significant reductions in the average daily drug dose, relative to continuous treatment. This benefit broadly increases with the AT threshold, with less drug required at higher tumor size thresholds, though the extent of this additional benefit varies between patients.

**Figure 9:**
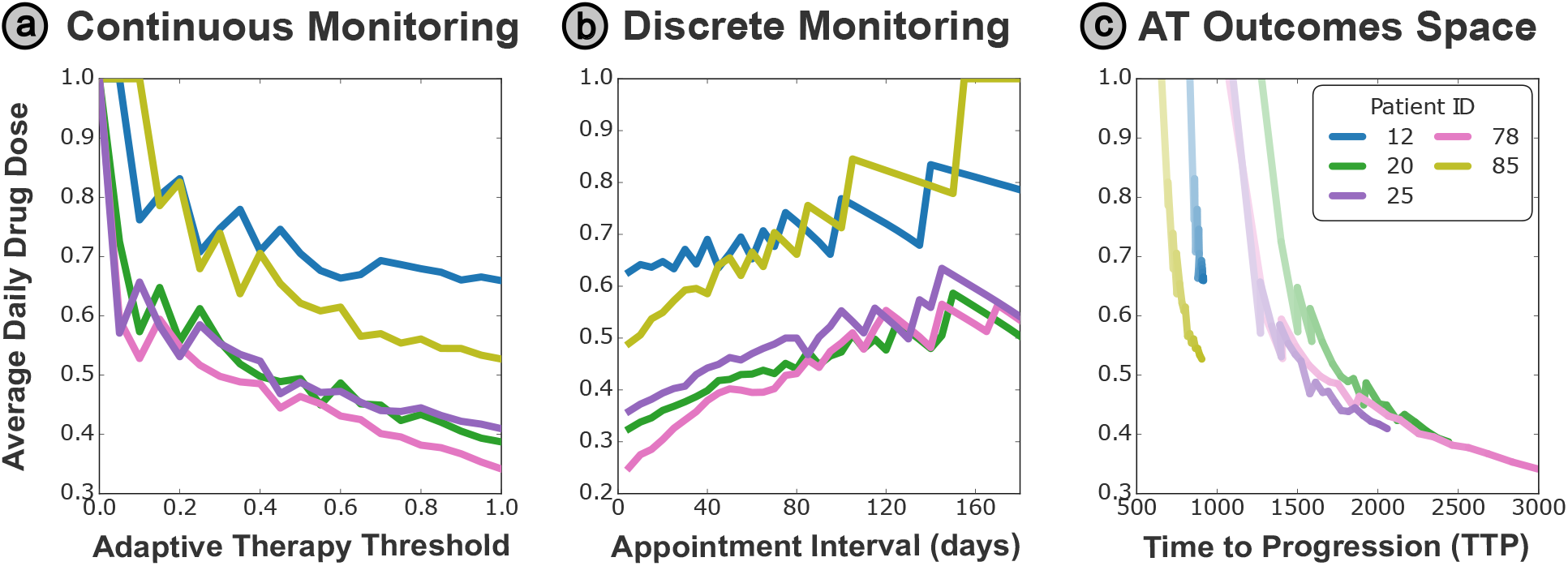
**(a)** Implementing adaptive schedules decreases the average daily drug dose significantly for all patients (relative to CT, with an average fractional dose of 1.0). Furthermore, this effect is still realized at very low treatment thresholds (where a threshold of 0.5 corresponds to withdrawing treatment when *N* (*t*) *<* 0.5*N*_0_). **(b)** This benefit is retained if the drug dosing is only updated at discrete appointment intervals, to better replicate the clinical reality of out-patient appointments. In this case we simulate the optimal strategy for each time interval between appointments, showing that all patients achieve significantly reduced average doses compared to CT at standard clinical appointment intervals (such as 60 days). **(c)** We can visualize the twin benefits of increased TTP and reduced average drug for a range of AT thresholds. We simulate treatment schedules with increasing adaptive thresholds (starting from CT where the average dose = 1.0), with the threshold size represented by the opacity of the color (where darker colors represent higher AT thresholds). We see the average drug decreasing and the TTP increasing (faster for some patients than others) as the AT threshold is increased from 0.

This trend is maintained when we considered clinically realistic implementations of AT, where PSA monitoring and treatment updating is limited to clinical appointments separated by a fixed time window. Shown in Figure 9b, we compute the average daily drug dose given under the optimal AT protocol at each appointment interval (where more frequent appointments allow higher AT thresholds and hence reduced average dosages). Again, all patients benefit from reduced intervals between appointments, with all patients benefiting from at least a 30% reduction in average drug dose at bimonthly appointments. It is also worth noting that all patients have a minimum interval where treatment breaks are viable; this is visible around 150 days for Patient 85, and they will simply be on drug treatment constantly when appointments are less frequent than this.

The combined benefit of both extended TTP and reduced average drug is plotted in Figure 9c, as the tumor size threshold is increased (where the intensity of the color denotes the tumor threshold, which darker colors representing higher thresholds). All patients lie on the upper limit of the plot for a zero size threshold (equivalent to CT as the tumor never reaches this size during treatment and hence no treatment holidays are given), but trend downwards and right as the threshold size is increased and treatment holidays are introduced for each patient with increasing frequency, under the optimal AT protocol for each patient. The gradient indicates the relative benefit each patient experiences from extended TTP and reduced average drug dose - the nearly vertical trajectories of Patients 12 and 85 indicates that they predominantly benefit from reduced average dosing. In contrast, the other patients mostly benefit from reduced dosing at long appointment intervals, but experience a much greater TTP benefit as the interval is further shortened.

